# Enhanced sampling of protein conformations in AlphaFold3 with repulsive bias in the diffusion generative model

**DOI:** 10.64898/2025.12.17.693105

**Authors:** Jun Ohnuki, Kei-ichi Okazaki

**Affiliations:** Research Center for Computational Science, Institute for Molecular Science, National Institutes of Natural Sciences, Okazaki, 444-8585, Japan; Graduate Institute for Advanced Studies, SOKENDAI, Okazaki, Aichi 444-8585, Japan

**Author notes:** Contributions J.O. and K.O. designed the research and wrote the paper. J.O. conducted the research and developed the methodology and the software. Corresponding author: Correspondence to: Jun Ohnuki & Kei-ichi Okazaki.

## Abstract

Conformational changes in proteins are vital to their function yet remain challenging for state-of-the-art artificial intelligence, such as AlphaFold3 (AF3), to predict. It has been observed that AF3 sometimes fails to capture ligand-induced conformational changes, even though it explicitly includes ligand molecules that induce such changes. To address this challenge, we develop an enhanced sampling scheme that leverages the diffusion-based generative model used in AF3 to predict protein structures. Interpreting the diffusion generative model as a stochastic sampling process analogous to molecular dynamics (MD) simulations, we introduce here a repulsive biasing potential between predicted structures to explore wider conformational space. We demonstrate that the developed model, AF3-ReD, successfully predicts multiple conformational states, including ligand-bound conformations of motor and transporter proteins that the original AF3 does not capture. Compared to another strategy based on multiple sequence alignment (MSA), AF3-ReD predicted intermediate conformations that are relatively closer to the stable states. Thus, AF3-ReD provides a promising approach for understanding dynamic conformational changes associated with ligand binding, including those induced by drug molecules.

## Introduction

Proteins change their conformation when they function. For example, motor proteins change their conformation in nucleotide-state-dependent manners to achieve the unidirectional motions^1–4^. Transporter proteins change conformation between inward-open and outward-open states to carry their substrates across the cell membrane^5,6^. Such conformational changes are vital to the function of these proteins, yet have been difficult for the state-of-the-art artificial intelligence (AI) for protein structure prediction, AlphaFold^7,8^, to predict. This posed a major challenge for AlphaFold2 (AF2) and led to the development of several pipelines to enable it to predict a wide range of conformations beyond a single conformational state^9,10^, including multiple-sequence-alignment (MSA)-related approaches^11–16^, template-related^17,18^ approaches, and integrations of AF2 with experimental information^19,20^ and other machine learning techniques^21,22^. The issue still persists in the latest version, AlphaFold3 (AF3), even though it explicitly includes ligand molecules that induce conformational changes^8^.

AF3 employs diffusion-based generative models^23–25^, a powerful class of generative AI methods, to predict protein structures. Diffusion-based generative models have been utilized to generate protein structures^8,26–30^, protein-ligand complexes^8,31^, sequences^32,33^, and to back-map atomic structures from coarse-grained models^34–36^. Additionally, they have been employed to learn coarse-grained force fields^37^ and to accelerate or extrapolate molecular dynamics (MD) simulations^38–40^. The diffusion model in AF3 is based on a score-based generative model using stochastic differential equations^24,25^, in which the score is optimized in training and corresponds to the learned gradient of the log probability density (see equation (1)). Assuming that the probability density follows the Boltzmann distribution, the score is equal to the gradient of an effective energy landscape^41^ (see equation (2)). Thus, the denoising process of the diffusion model in AF3 can be seen as simulating the formation of a protein conformation along an energy landscape (Fig. 1a). Therefore, the limited prediction ability of AF3 to a single conformational state is somewhat similar to the sampling problem of rare events in MD simulations. In MD simulations, a system is typically trapped in a metastable state and unable to visit other metastable states due to the timescale of barrier crossing events. Various enhanced sampling techniques have been developed to overcome this problem, including umbrella sampling^42,43^, replica exchange^44–46^, steered MD^47–50^, transition path sampling^51–54^, accelerated MD^15,55–57^, and metadynamics^58,59^. Some of these methods apply an artificial force to enhance sampling of a wider region in conformational space. Given that AlphaFold has effectively learned an energy function^60^ that has multiple basins, which describes conformational changes^61,62^ and is indicated by the distance probability distribution from AlphaFold^63,64^, such enhanced sampling techniques would also help AF3 overcome its limitations.

**Fig. 1:**
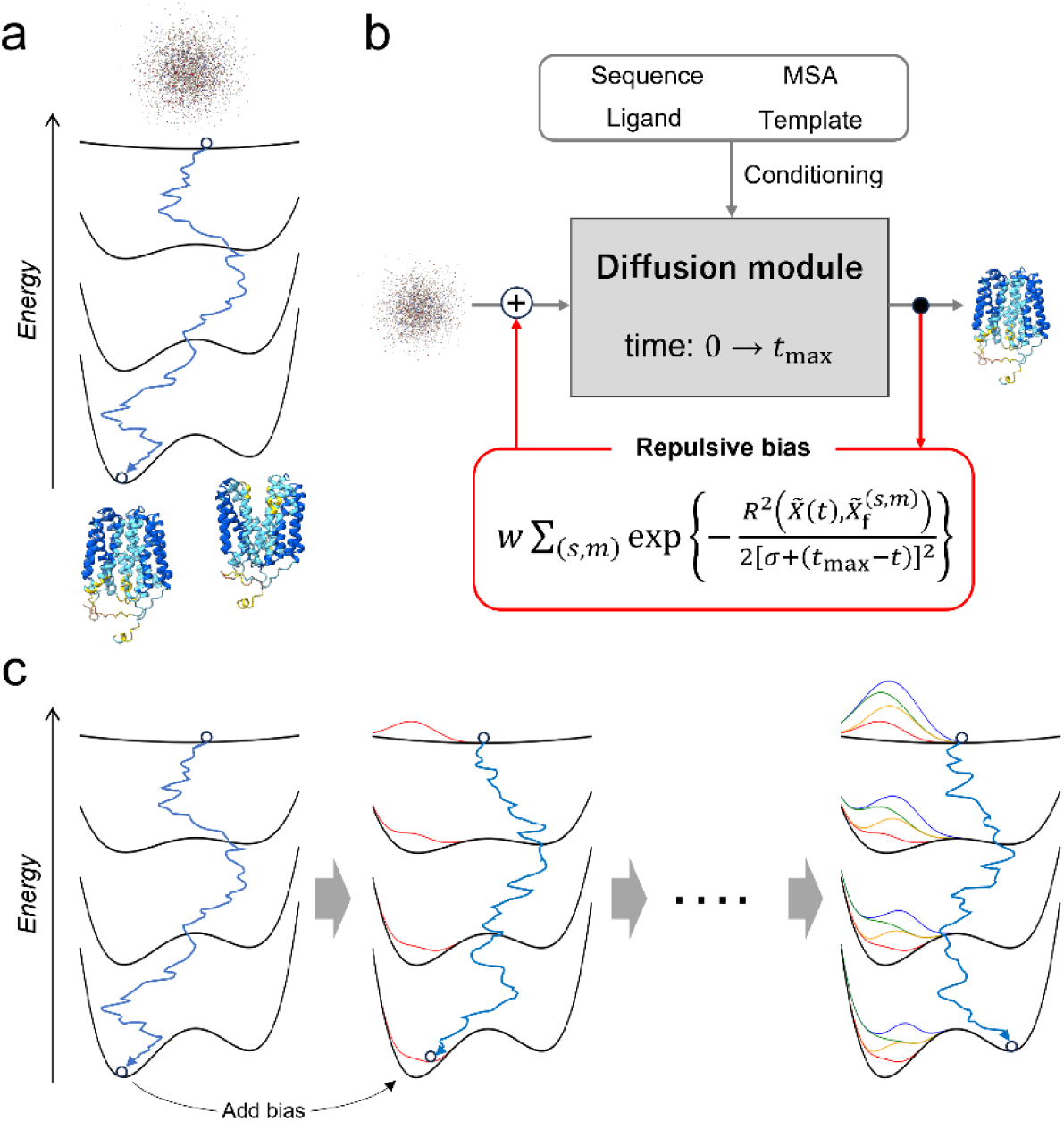
Repulsive biasing potential introduced in the denoising process of the AlphaFold3 diffusion model (AF3-ReD) **a** Stochastic sampling of protein conformations with the denoising process of the AlphaFold3 diffusion model. **b** Architecture of AlphaFold3 diffusion model with repulsive biasing potential. The red box and arrows indicate the introduced biasing potential. **c** Stochastic conformational sampling with the denoising process in the presence of repulsive bias.

In this work, we introduce AlphaFold3 with a repulsive biasing potential applied to the diffusion model (AF3-ReD) to enhance conformational sampling of proteins. Inspired by the metadynamics method^58,59^, the structure generation in AF3-ReD is guided by repulsive biasing forces from previously predicted structures during the denoising process, enabling the diffusion model to avoid generating redundant structures within the same conformational basin. We also implemented one of the most widely used MSA-related methods, MSA subsampling^11^, in AF3 and compared its performance to that of AF3-ReD. Application of AF3-ReD to motor and transporter proteins in ligand-bound conditions demonstrated that it successfully predicts multiple conformational states, including ligand-bound conformations typically not captured by the default AF3 or AF3 with MSA subsampling. AF3-ReD achieved conformational diversity without large deviation from stable states, representing plausible intermediate conformations. It can also be applied to multimer complexes, although capturing large-scale conformational changes in protein complexes remains challenging. The enhanced sampling scheme of AF3-ReD not only provides a foundation for understanding dynamic conformational changes but also offers insights for improving other diffusion-based generative models for protein structures.

## Results

### Repulsive biasing potential in AlphaFold3 diffusion model (AF3-ReD)

AF3 takes protein and DNA/RNA sequences, ligand chemical information, and template structures as input, and computes embeddings from these inputs as well as MSAs from the sequences. The resulting embeddings are then used to condition the score function, which corresponds to the learned gradient of the log probability density:

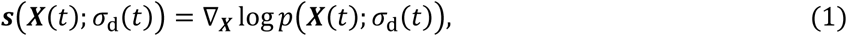

where ***X***(*t*) is protein structure coordinates at time *t* and *p*(***X***; σ_d_) is the probability density at a given noise level σ_d_(*t*) in the denoising process. Assuming that the probability density follows the Boltzmann distribution, the score is equal to the gradient of an effective energy landscape^41^:

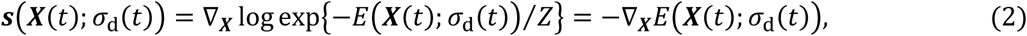

where *Z* is the normalizing constant (i.e., partition function) unrelated to the score calculation. In this way, the energy landscape is constructed from the input information. The conditioned score function is applied to the denoising process in the diffusion module of AF3, where initial random noise is gradually denoised according to the time-dependent energy landscape to yield a final structure (Fig. 1a). Although the AF3 diffusion model can sample protein conformations following the energy landscape during the denoising process, imbalances in the learned distributions arising from the training data make it difficult to sample alternative conformational states.

To avoid predicting similar structures within the same conformational state, in analogy to the history-dependent repulsive biasing potential used in widely used metadynamics simulations, we introduce a Gaussian repulsive potential during the denoising process of the AF3 diffusion model (Fig. 1b and 1c). Root-mean-squared deviation (RMSD) is used as a collective variable for the repulsive potential. For the *m*-th prediction model with the *s*-th random seed (*m* =1 … 5 since typically five models are generated per seed in AF3), we denote the Cα-atom coordinates of a predicted structure as 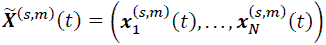, where 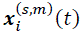 represents the position of the *i*-th Cα atom at time *t* and *N* is the number of Cα atoms. For conciseness, the set of the Cα coordinates of the current structure in the diffusion process is denoted as 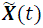 without the superscripts. As AF3 sequentially performs the denoising process for each random seed, RMSD is calculated between the current structure and all previously predicted structures:

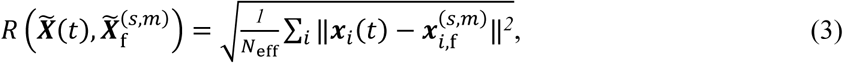

where *N*_eff_ is the number of atoms used in the RMSD calculation (see Methods for details), and 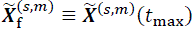 is the Cα coordinates of a previously predicted model (the *m*-th prediction with the *s*-th random seed) with the final time *t*_max_ = 160 in AF3, meaning a final denoised structure. The Gaussian repulsive potential is defined using the RMSD as a collective variable:

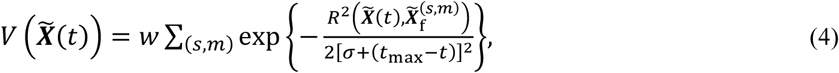

where *w* and *σ* are the strength and width of the Gaussian potential, respectively. To avoid applying the biasing potential to alternative conformations, the sum is taken only over structures that are close to the ones from the first random seed (see Methods for details).

Adding the bias potential makes the denoising process experience repulsive forces from previously predicted structures. Because the denoising process lacks constraints between adjacent atoms, such as covalent bonds, raw repulsive forces are not effective at facilitating concerted motions required for transitions to alternative conformations. Thus, the repulsive force acting on the *i*-th Cα atom is smoothed over the adjacent 10 Cα atoms on each side:

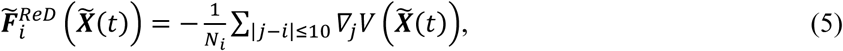

where *N*_*i*_ represents the number of Cα atoms used for smoothing. In addition, the force acting on each Cα atom is applied to all atoms within the residue to which that Cα atom belongs. The obtained force is referred to as 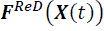, and is added during the positional update in the denoising process as follows:

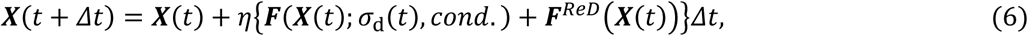

where ***F***(⋅) represents the effective force of the original AF3, which is proportional to the score function conditioned on the embeddings (*cond*.) at a given noise level σ_d_(*t*), and *η* is a step-scale parameter, set to 1.5 in AF3. Stochastic sampling with the denoising process proceeds by alternately repeating this position update and a noise-adding step^8,25^. Note that in AF3, Δ*t* is defined as negative due to the reverse diffusion process, but we set Δ*t* to positive for clarity, so that the forces act in the direction of decreasing energy, as in standard MD simulations.

It is worth noting that our developed AF3-ReD requires no retraining and just adds the biasing potential to the energy landscape trained by the original AF3. A recent study showed that the introduction of repulsive biasing forces into the diffusion model promotes diversity in image generation and conformation generation of small molecules without further training^65^. In the following sections, we investigated whether AF3-ReD enhances sampling of alternative conformations for target proteins.

### Application of AF3-ReD to the β subunit of F_1_-ATPase

As a target of our new approach, AF3-ReD, we look into the β subunit of the rotary motor F_1_-ATPase from thermophilic *Bacillus* (TF_1_β), which binds ATP and changes its conformation from the open to the closed conformation (Fig. 2a). The three β subunits, the three non-catalytic α subunits, and the central shaft γ subunit constitute the F_1_-ATPase complex, one of the best-studied biomolecular motors^49,66–74^. The conformational change of the β subunit is a major driving force of the unidirectional rotation of the γ subunit. The nuclear magnetic resonance (NMR) experiment previously showed that the monomeric TF_1_β by itself changes its conformation upon ATP binding^75^.

**Fig. 2:**
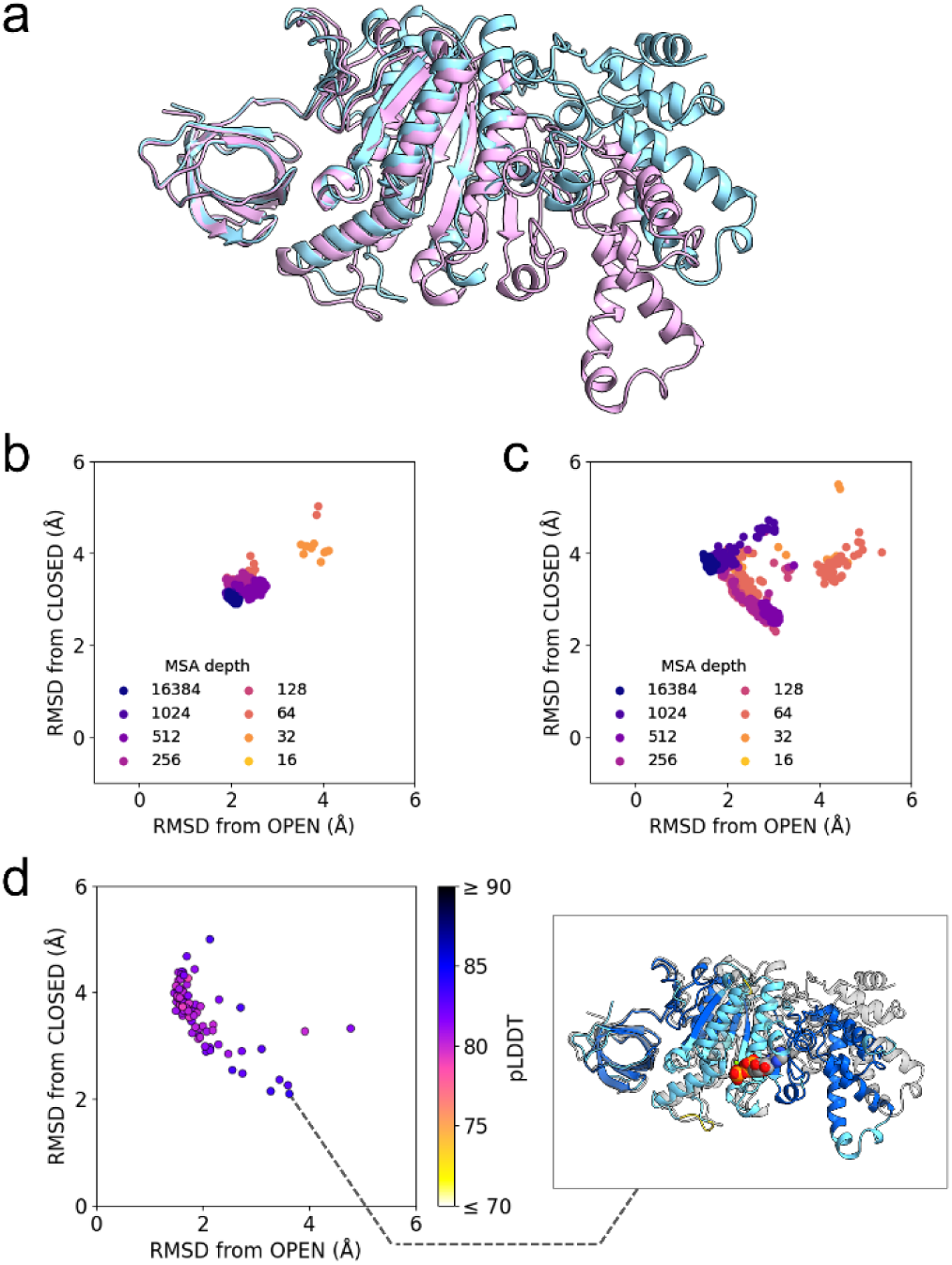
Structure prediction of F_1_-ATPase β subunit with AF3-MSA-subsampling and AF3-ReD. **a** Structures of the β subunit of F_1_-ATPase from thermophilic *Bacillus* (TF_1_β) in the open (cyan) and closed (pink) states. Both TF_1_β structures were extracted from the F_1_-ATPase complex determined by cryo-electron microscopy (PDB ID: 7XKH). **b, c** Root-mean-square deviations (RMSDs) of the structures predicted with MSA-subsampled AlphaFold3 for the apo TF_1_β (b) and the ATP-bound TF_1_β (c). RMSDs were calculated using the Cα atoms of residues 1–470 with respect to the open and closed cryo-EM structures. **d** RMSDs of the structures predicted with AF3-ReD (w = 90, σ = 2 Å) for the ATP-bound TF_1_β with respect to the open and closed cryo-EM structures. Points are colored according to the pLDDT score averaged over all atoms. A predicted closed conformation is also shown, where amino acid residues are colored according to their pLDDT score (blue: pLDDT ≥ 90; cyan: 70 ≤ pLDDT < 90; yellow: 50 ≤ pLDDT < 70; orange: pLDDT < 50).

First, the default AF3 predicted the TF_1_β conformation solely in the open state under both apo and ATP-bound conditions (blue points with the MSA depth 16,384 in Fig. 2b and 2c, respectively), a typical limitation of AF3. Then, we implemented the previously proposed shallow MSA subsampling approach^11^ in AF3 by treating the MSA depth as a variable, serving as a baseline for our new method (see Methods for details). In the apo condition, the AF3 MSA subsampling basically predicted the open conformations (Fig. 2b). This was also observed with the AF2 MSA subsampling, which does not consider ligands (Supplementary Fig. 1). In the ATP-bound condition, as the MSA depth was reduced, the AF3 MSA subsampling extended the conformational distribution and predicted closed conformations with root-mean-squared deviations (RMSDs) to the experimental closed structure^73^ near 2 Å (Fig. 2c).

As with the AF3 MSA subsampling, our AF3-ReD successfully predicted a wide range of conformations with bound ATP, from the open to the closed state (Fig. 2d), including intermediate conformations. This result indicates that our AF3-ReD approach with the repulsive biasing force is effective at sampling diverse conformations beyond the most populated open conformation. Compared to the AF3 MSA subsampling, AF3-ReD predicted conformations that are close to either the open or the closed experimental conformations. On the contrary, the AF3 MSA subsampling occasionally predicted conformations that are far apart (RMSD > 3 Å) from both the open and closed experimental conformations. It should also be noted that although AF3-ReD introduces additional biasing forces in the diffusion process, its computational cost remained almost unchanged. Generation of 100 protein structures (excluding MSA generation) for the TF_1_β took 12 min 13 s with AF3 and 12 min 24 s with AF3-ReD using an AMD EPYC 7763 (16 cores) and an NVIDIA A100 GPU.

### Application of AF3-ReD to transporter proteins

We next apply AF3-ReD to transporter proteins that change conformation between outward-open and inward-open states to carry their substrates across the membrane. They typically have an intermediate occluded state in which the binding site is occupied with the substrate and not accessible from either side of the membrane. Previous studies using AF2 showed that it predicts either the inward or outward state with the default MSA depth, and as the MSA depth decreases, predicted conformations extend to the occluded and finally reach the alternative conformation^11,16^. The major limitation of AF2 here is that the occluded conformation lacks the bound substrate, a problem addressed by AF3.

First, we investigated the oxalate transporter OxlT. The apo outward-open and oxalate-bound occluded structures were previously solved by X-ray crystallography^76^ (left and center panels of Fig. 3a). The Gaussian-accelerated MD simulation, one of the enhanced-sampling methods with external biasing potentials, revealed the remaining inward-open conformation (right panel of Fig. 3a)^15^. AF3 with the default MSA depth in apo or oxalate-bound conditions already predicted both the inward-open and outward-open conformations (Fig. 3b and 3c). As the MSA depth decreases, the inward-open and outward-open conformations were slightly extended. However, the prediction does not overlap with either the occluded crystal structure or its corresponding MD ensemble (green and gray points in Fig. 3b and 3c, respectively), indicating that the AF3 MSA subsampling is limited in predicting the substrate-bound occluded conformation of OxlT. In contrast, AF3-ReD efficiently predicted the occluded conformation with oxalate bound (Fig. 3d). The predicted conformations overlapped with the MD-sampled occluded region, demonstrating AF3-ReD’s improved ability to sample the substrate-bound occluded state.

**Fig. 3:**
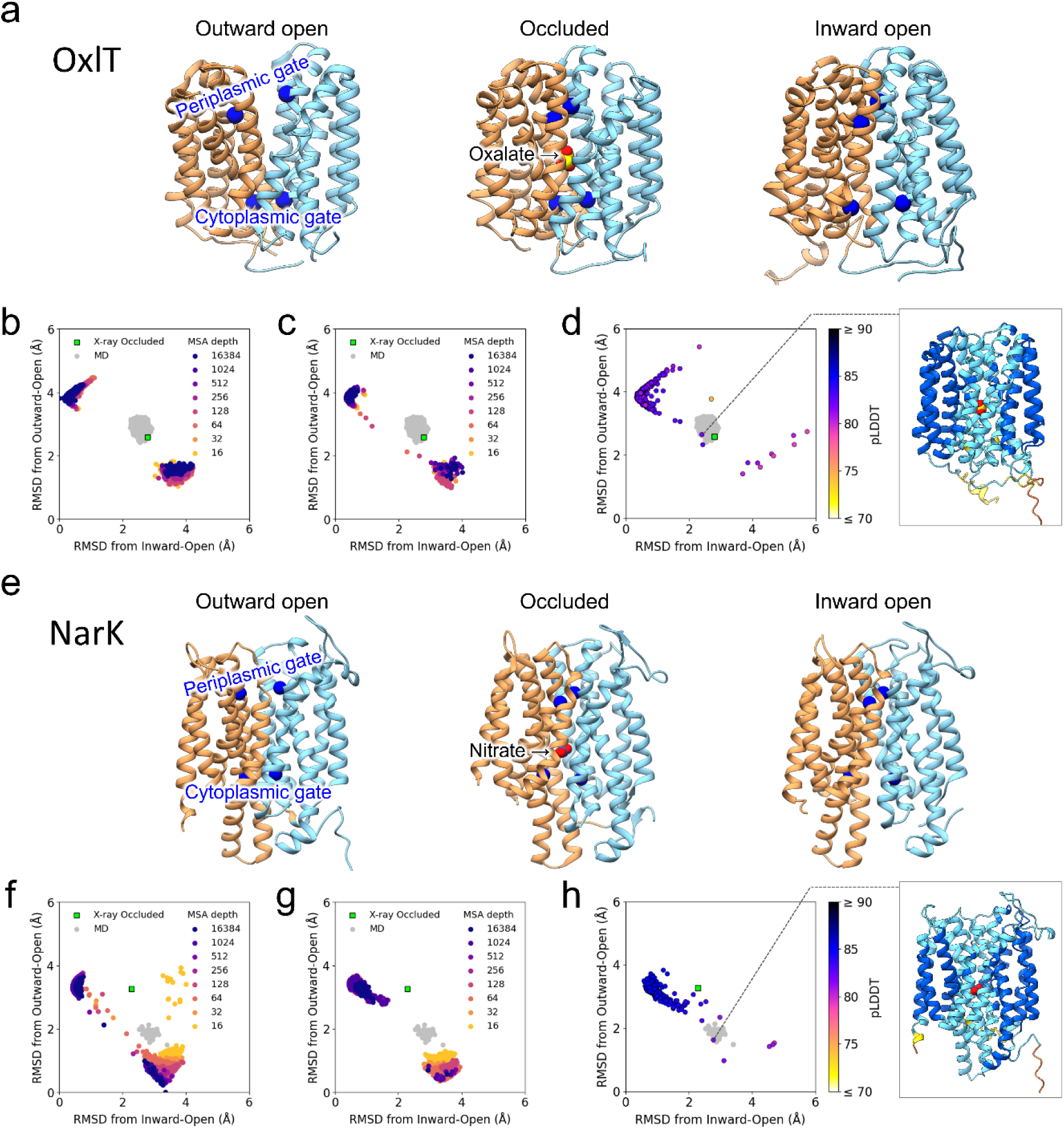
Structure prediction of transporter proteins with AF3-MSA-subsampling and AF3-ReD. **a** Structures of OxlT in the outward-open, occluded, and inward-open states. The outward-open and occluded structures were previously obtained by X-ray crystallography (PDB ID: 8HPJ and 8HPK, respectively). The inward-open structure was previously predicted by our molecular dynamics study. **b, c** RMSDs of the structures predicted with MSA-subsampled AlphaFold3 for the apo OxlT (b) and for the oxalate-bound OxlT (c). RMSDs were calculated using the Cα atoms of residues 11–196 and 204–405 with respect to the outward-open crystal structure and the inward-open structure predicted with the default AF3. For the latter, the prediction with the highest pLDDT for the apo OxlT was used as the reference. The occluded crystal structure (green) and the MD simulation structures initiated from it (gray) are also plotted. **d** RMSDs of the structures predicted with AF3-ReD (w = 10, σ = 1 Å) for the oxalate-bound OxlT. A predicted occluded conformation is also shown, where amino acid residues are colored according to their pLDDT score, as shown in Fig. 2d. **e** Structures of NarK. The inward-open and occluded structures were previously obtained by X-ray crystallography (PDB ID: 4U4V and 4U4W, respectively). The outward-open structure was previously predicted in our previous study using AF2. **f, g** RMSDs of the structures predicted with MSA-subsampled AlphaFold3 for the apo NarK (f) and for the nitrate-bound NarK (g). RMSDs were calculated using the Cα atoms of residues 14–239, 252–342 and 344–457 with respect to the inward-open crystal structure (PDB ID: 4U4V) and the outward-open structure predicted with the default AF3. For the latter, the prediction with the highest pLDDT for the apo NarK was used as the reference. The occluded structures from X-ray crystallography (green, PDB ID: 4U4W) and from our previous MD simulation (gray) are also plotted. **h** RMSDs of the structures predicted with AF3-ReD (w = 8, σ = 1 Å) for the nitrate-bound NarK. A predicted occluded conformation is also shown.

Next, we applied our AF3-ReD to the nitrate transporter NarK. The apo/nitrate-bound inward-open and nitrate-bound occluded structures were previously solved by X-ray crystallography^77^ (Fig.3e). We previously showed that decreasing the MSA depth in AF2 successfully predicted broad conformations of NarK, including the experimentally unresolved outward-open state^16^ (Supplementary Fig. 3). Furthermore, MD simulations initiated from an AF2-predicted intermediate conformation found the occluded conformation with the cytoplasmic gate more closed than the occluded crystal structure, which is closer to the outward-open state (gray points in Fig. 3f and 3g). As already seen with OxlT, using the default MSA depth, AF3 already predicted both inward-open and outward-open conformations in the apo or nitrate-bound conditions (Fig. 3f and 3g), and a reduction in the MSA depth further broadened the distribution. However, we observed no overlap between the AF3 MSA subsampling distribution and either the occluded crystal or the MD-generated structures. AF3-ReD, in contrast, predicted the occluded state close to the crystal or MD structures (Fig. 3h). These results indicate that AF3-ReD is more efficient in sampling substrate-bound occluded conformations of transporter proteins than the AF3 MSA subsampling.

### Sampling coverage and diversity

Thus far, our results demonstrate that AF3-ReD enhances conformational sampling efficiency in the presence of substrate molecules. To obtain quantitative insights, we compared the sampling coverage and diversity of AF3-ReD with those obtained from the MSA-subsampling method and examined their dependence on model parameters. The sampling coverage was defined as the fraction of predicted structures whose RMSD from either of the two reference structures (the open/closed states for the TF_1_β, and the inward/outward-open states for the transporter proteins) was smaller than a given threshold (Fig. 4a). The coverage at an RMSD threshold of 3 Å was used to compare AF3-ReD with the AF3 MSA subsampling. The sampling diversity was defined as the exponential of the Shannon entropy^78,79^ estimated from the probability density of RMSDs with respect to the reference structures, and normalized by that obtained from the default AF3 (see Methods for details).

**Fig. 4:**
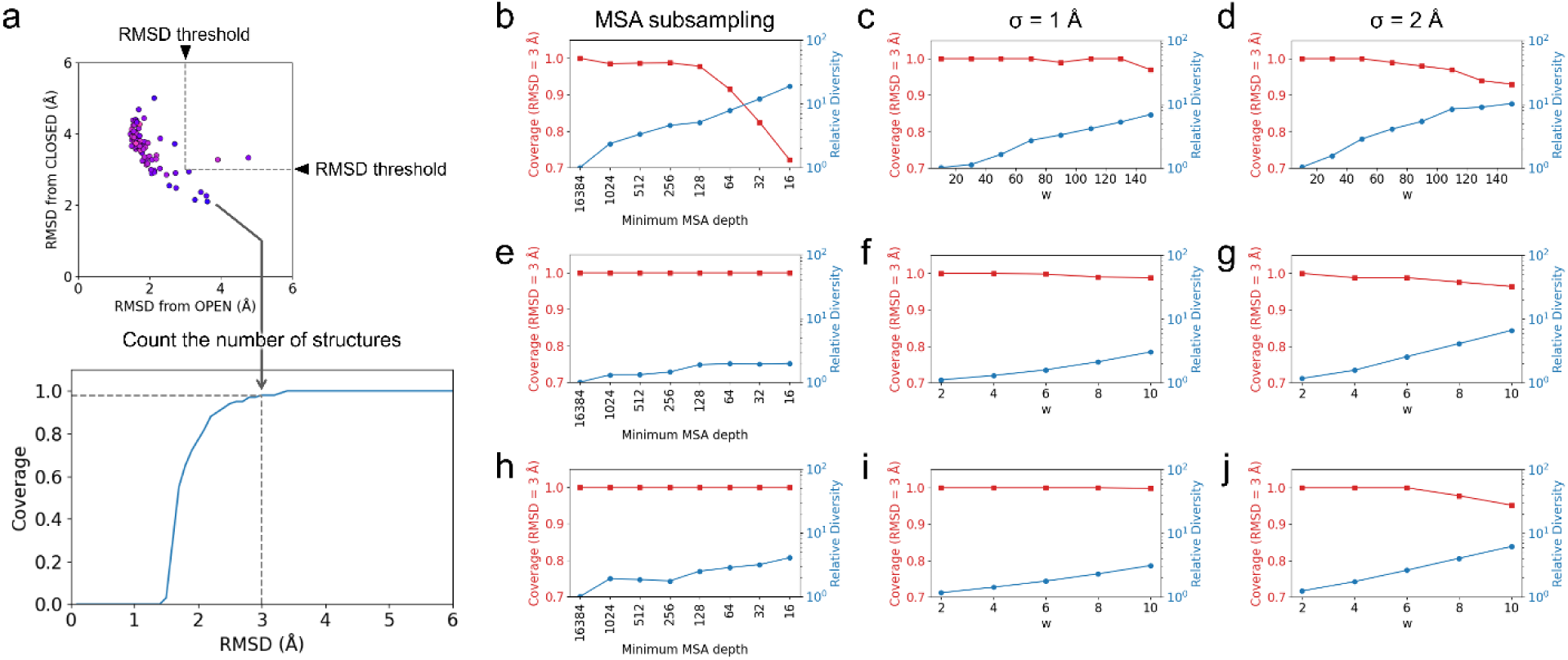
Sampling coverage and diversity of AF3-MSA-subsampling and AF3-ReD. **a** An illustrative example of sampling coverage analysis. The coverage is defined as the proportion of structures whose RMSD from either of the two reference structures is below a given threshold. **b-j** Sampling coverage and diversity for TF_1_β (b-d), OxlT (e-g), and NarK (h-j). For each protein, results are obtained using different methods and parameter settings: AF3 with MSA subsampling (b, e, h), AF3-ReD with σ = 1 Å (c, f, i), and AF3-ReD with σ = 2 Å (d, g, j).

The sampling coverage and diversity of the TF_1_β in the holo condition are shown in Fig.4b for the MSA subsampling and in Fig. 4c and 4d for AF3-ReD. For the MSA subsampling, we used all structures obtained from MSA depths above a given depth; for example, the sampling coverage and diversity for the MSA depth of 16 were calculated using all the depths tested here. Decreasing MSA depth for the holo TF_1_β increased the diversity but markedly worsened the coverage, which is consistent with the observation in Fig. 2c. In contrast, AF3-ReD with σ = 1 Å demonstrates that increasing the weight of the bias potential leads to a higher diversity while maintaining a relatively high level of coverage (Fig. 4c). A larger σ leads to a further increase in the diversity, accompanied by a slight decrease in the coverage (Fig. 4d). The coverage and the relative diversity with σ = 2 Å and w = 90 (0.98 and 5.4, respectively), as used in Fig. 2d, are comparable to the values obtained with MSA subsampling at a depth of 128 (0.98 and 5.1, respectively). It should be noted, however, that the MSA-subsampling result shown here was obtained using all predicted structures generated with depths of 128 or higher (i.e., from 128 to 16,384), and that when only the predictions with a depth of 128 are used, the relative diversity decreases to 2.6. These results indicate that AF3-ReD can achieve diverse conformational sampling with fewer generated structures.

Figures 4e to 4j show the sampling coverage and diversity of the transporter proteins obtained by the AF3 MSA subsampling and our AF3-ReD. Compared to the TF_1_β, the relative diversity in the MSA subsampling increases only slightly as the MSA depth decreases (Fig. 4e for OxlT and Fig. 4h for NarK). In contrast, the biasing potential of AF3-ReD increases the diversity as the weight *w* and width *σ* increase, achieving higher diversity than that obtained with the AF3 MSA subsampling while maintaining high coverage values (Fig. 4f, g for OxlT and Fig. 4i, j for NarK). These results further support the conclusion that AF3-ReD efficiently enhances conformational sampling compared to the AF3 MSA subsampling.

### Application of AF3-ReD to the E3 ubiquitin ligase complex

We further tested our AF3-ReD with a larger multi-subunit system, the E3 ubiquitin ligase complex. The E3 ubiquitin ligase complex, composed of cereblon (CRBN), the DNA damage-binding protein 1 (DDB1), Cullin-4, and the RING finger protein ROC1, transfers ubiquitin to substrate proteins. A previous cryo–electron microscopy (cryo-EM) study solved part of the complex, CRBN-DDB1, showing the open CRBN in the apo state and the closed CRBN in the mezigdomide-bound state^80^. The observed structures reveal a substantial conformational change (RMSD > 10 Å) in the complex (Fig. 5a). In the original AF3 paper, Abramson et al. reported that AF3 is unable to capture the conformational state of the CRBN-DDB1 complex, as the closed state is predicted even in the apo condition^8^. Thus, the CRBN–DDB1 complex serves as a challenging target for testing the capability of AF3-ReD.

**Fig. 5:**
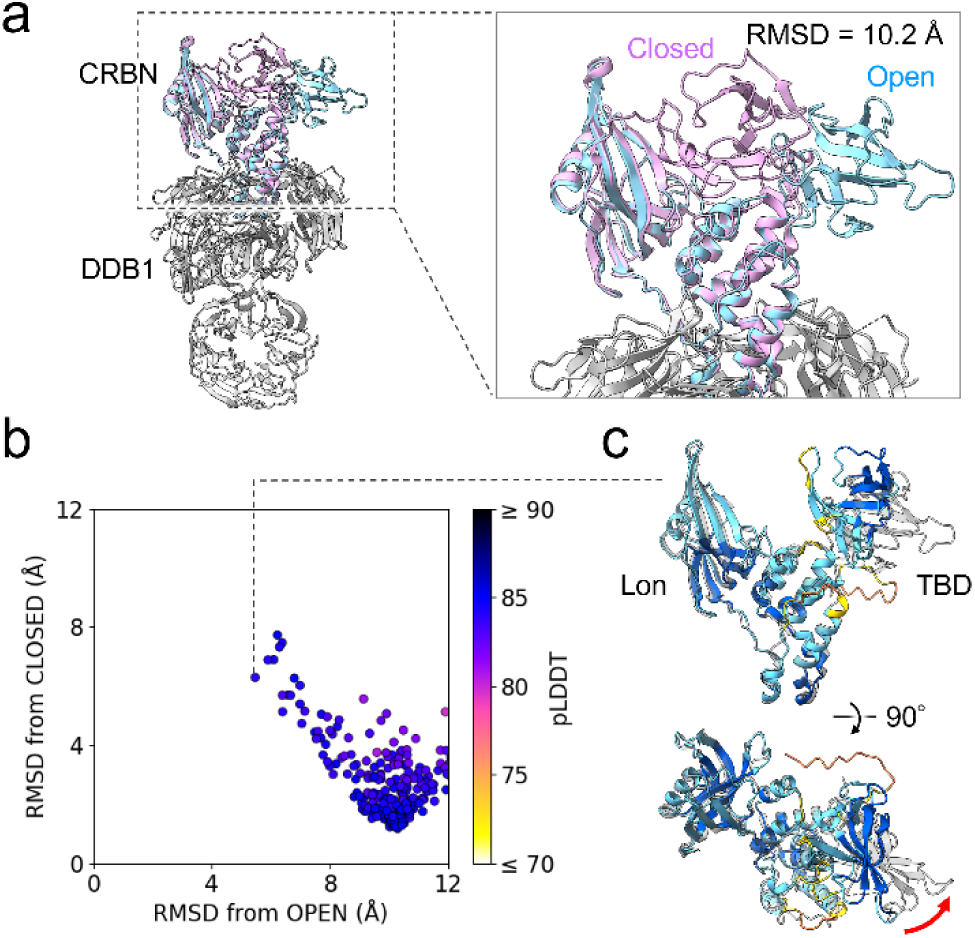
Structure prediction of the E3 ubiquitin ligase complex with AF3-ReD. **a** Structures of the E3 ubiquitin ligase complex CRBN-DDB1 in the open and closed states (PDB ID: 8CVP and 8D7U, respectively). **b** RMSDs of the structures predicted with AF3-ReD (w = 100, σ = 2 Å) for the apo CRBN-DDB1. RMSDs were calculated using the Cα atoms of CRBN residues 64–341 and 358–425 with respect to the open and closed cryo-EM structures. **c** A partially-open conformation predicted with AF3-ReD. Amino acid residues are colored according to their pLDDT score, as shown in Fig. 2d. The red arrow indicates the rotation required to complete the transition to the fully-open conformation (transparent gray).

We here predicted the structures of the CRBN–DDB1 with the N-terminal region of CRBN (residues 1 to 63) deleted, because this region is reported to stabilize the closed conformation^80^. First, the default AF3 predicted the apo CRBN–DDB1 complex exclusively in the closed conformation, even after deletion of the N-terminal region (Supplementary Fig. 4a). AF3 with MSA subsampling produced structures closer to the open state when the MSA depth was set to 256. However, these structures exhibited very low pLDDT scores (Supplementary Fig. 4b), and the relative orientation between CRBN and DDB1 deviated substantially from the experimental structure even though the BPB domain of DDB1 is known to be flexible^80^ (Supplementary Fig. 4c), highlighting the difficulty of predicting complex structures via the AF3 MSA subsampling. In contrast, AF3-ReD extended the conformational distribution toward the open state while maintaining high pLDDT scores (Fig. 5b). Nevertheless, the predictions did not achieve an RMSD below 5 Å from the experimental open structure. Comparison with the experimental structure revealed that, although the two CRBN domains (Lon and TBD) were sufficiently separated, AF3-ReD failed to capture the additional rotational motion of the TBD (Fig. 5c). Taken together, these results indicate that predicting such large-scale conformational changes in protein complexes remains challenging even with the AF3 MSA subsampling or AF3-ReD, underscoring the need for further methodological improvements.

## Discussion

We here developed AF3-ReD, where the repulsive biasing potential, inspired by the metadynamics technique^58,59^, was introduced to the AF3 diffusion model to enhance sampling of protein conformations. Applications of AF3-ReD to the β subunit of F₁-ATPase and two transporter proteins demonstrated its ability to predict substrate-induced conformational changes of these proteins, which was not captured by the original AF3. While AF3 with MSA subsampling also succeeded in predicting the substrate-bound conformation of the F_1_ β subunit, it failed to do so for the transporters. These results indicate that the effectiveness of MSA subsampling strongly depends on the protein system. This is likely due to differences in the underlying biases in MSAs that prevent the default AF3 from exploring alternative conformations. For the F_1_ β subunit, the increase in diversity upon decreasing the MSA depth indicates that the coevolutionary information in the MSA, used to condition the score function of the AF3 diffusion model, favors a specific conformational state^15^. In contrast, the weaker MSA-depth dependence of sampling diversity for the transporter proteins suggests that the difficulty in sampling arises from other sources inherent in the AF3 diffusion model, such as biases in the AF3 training data and insufficient sampling time at high noise levels due to the noise schedule. Unlike the MSA subsampling, AF3-ReD can mitigate both coevolution-derived and diffusion-model-derived sampling difficulties, as it directly modulates the energy landscape determined by the training data and the MSA-derived coevolution. Nevertheless, even AF3-ReD has yet to capture the large-scale conformational change observed in the E3 ubiquitin ligase complex, indicating that further improvements in conformational sampling are needed. Such improvements may include combining AF3-ReD with other approaches, such as MSA subsampling, redefining the biasing potentials in AF3-ReD using different CVs, or refining the AF3 energy landscape itself (i.e., the neural network parameters).

Although AlphaFold generates only static structural information, integrating its predictions with MD simulations can provide deeper insights into conformational dynamics and energetics^16,81,82^. We previously demonstrated that AF2 with MSA subsampling can generate transition-state-like conformations of NarK that lie between the stable conformational states^16^. Such intermediate conformations were also observed in predictions by AF3-ReD. These structures can serve as starting points for rare-event MD sampling, such as Markov state model^83,84^ or transition path sampling^51–54^, thereby enabling efficient characterization of conformational dynamics and thermodynamics in proteins. Besides, MD simulations themselves may even be replaceable by generative AI methods. For example, recently developed BioEmu generates a conformational ensemble of proteins through a diffusion-based generative model that learned MD simulations and experimental stability data^30^. Therefore, generative AI methods and MD simulations can mutually reinforce each other, and such bidirectional integrations should provide fruitful insights into dynamic conformational changes associated with ligand binding, such as drug molecules.

Diffusion models are powerful AI frameworks that have been widely applied to the prediction and design of biomolecules, as well as to the generation of ligands and drugs that bind to them^85^. However, the quality of the generated samples often comes at the expense of their diversity^86^, as also observed in AF3. To address this issue, AF3-ReD introduces repulsion between sampled structures during the denoising process. The utility of such repulsive mechanisms in diffusion models has also been demonstrated in image generation and small-molecule conformation generation^65,87^. In AF3-ReD, the repulsive biasing potential is defined simply by the structural similarity (i.e., RMSD) between sampled structures, which facilitates diverse conformational sampling without retraining. This strategy could be further extended to other diffusion-based generative models for designing protein structures, potentially expanding their accessible design space.

## Methods

### Repulsive biasing potential in AlphaFold3 diffusion model (AF3-ReD)

The repulsive biasing potential was implemented in AF3 (code version as of Mar 17, 2025, commit ID: 8151373). Before calculating RMSDs according to equation (3), all previously predicted structures were superimposed onto the current structure during the denoising process of the diffusion module using the Kabsch algorithm^88,89^. In the case of protein complexes, either the entire complex or only specific subunits were used for the superposition; for example, in the CRBN–DDB1 complex, only CRBN was aligned here. For the superimposition and RMSD calculation, we define the atom set *A,* which contains the indices of Cα atoms whose pLDDT scores predicted with the first random seed are greater than 60:

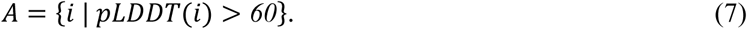

If there are multiple prediction models for the first random seed, the pLDDT scores are averaged over the models. We superimposed structures and calculated RMSDs using atoms included in the subset *A*. This pLDDT-based selection helps exclude disordered regions, which typically have low pLDDT scores, from the RMSD calculation.

When calculating the repulsive biasing potential, the sum in equation (4) is taken over the set *S* of previously predicted structures for which the RMSD from at least one model with the first random seed is less than 0.5 Å:

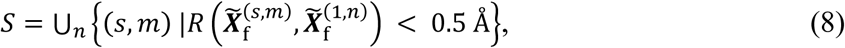

where *s* and *m* represent the indices for the random seed and prediction model, respectively. This restriction prevents us from applying the biasing potential to alternative conformations. The gradient of the biasing potential was added to the denoising process, as described in equation (6), except in the final stage (*t*_max_ − *t* < 2.0) where no biasing force is applied to refine the structures.

### MSA subsampling in AlphaFold3

In AF3 (code version as of Mar 17, 2025, commit ID: 8151373), the maximum number of homologous sequences in the multiple sequence alignment (MSA), referred to as the MSA depth, is hard-coded with a default value of 16,384. To enable control of this parameter, we modified the AF3 source code so that the MSA depth could be specified as a variable. For the proteins studied here, the MSA depth was varied in the range from 16,384 (default) to 16.

### Structure prediction of proteins

The amino acid sequences of the proteins used in this study (UniProt IDs) were as follows: P07677 for the β subunit of F_1_-ATPase from thermophilic *Bacillus* (TF_1_β), Q51330 for OxlT, P10903 for NarK, Q96SW2 for CRBN, and Q16531 (excluding the initiator methionine) for DDB1. For OxlT and NarK, the MSAs generated in our previous studies^15,16^ were used. The N-terminal region of CRBN (residues 1–63) was truncated to facilitate sampling of the open conformation. The MSA depth in the AF3 MSA subsampling, as well as the strength and width of the Gaussian potential in AF3-ReD, were varied to investigate the effects of these parameters on conformational sampling. For each random seed, AF3 generated five model structures, and the total number of predicted structures for each parameter setting was as follows: 100 structures (20 seeds × 5 models) for TF_1_β (apo and ATP-bound conditions), 500 (100 × 5) for OxlT (apo and oxalate-bound conditions), 500 (100 × 5) for NarK (apo and nitrate-bound conditions), and 300 (60 × 5) for the CRBN–DDB1 complex (apo condition). For comparison, structure prediction with AlphaFold2 was performed using ColabFold^90^ version 1.5.2, which is compatible with AlphaFold 2.3. No templates were used in any of the structure predictions. Molecular graphics were performed with UCSF Chimera^91^ and ChimeraX^92^, and RMSD analysis was performed with MDAnalysis^93,94^.

### Sampling coverage and diversity

The sampling coverage was calculated to evaluate the ability of the AF3 MSA subsampling and AF3-ReD to predict conformations close to stable states. We first counted the number of predicted structures whose RMSD from either of the two reference stable structures was smaller than a given threshold (Fig. 4a). As the references, the open and closed states were used for TF_1_β, and the inward- and outward-open states were used for the transporter proteins. The coverage was then obtained by dividing that number by the total number of predictions, yielding a value between 0 and 1. The coverage at an RMSD threshold of 3 Å was used to compare AF3-ReD with the AF3 MSA subsampling.

The sampling diversity was calculated to evaluate the ability of the AF3 MSA subsampling and AF3-ReD to explore a broad range of conformations. It was defined as the exponential of the Shannon entropy *S*^78,79^:

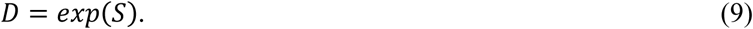

The Shannon entropy was estimated from the probability density of RMSDs with respect to the reference structures:

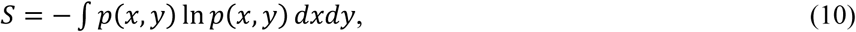

where *x* and *y* are the RMSDs from each of the two reference structures, and *p*(*x*, *y*) is the probability density. *D* has a desirable property as a measure of diversity: *D* doubles when the range of an equiprobable distribution doubles. The relative diversity was then obtained by dividing *D* by that from the default AF3, and compared between AF3-ReD and the AF3 MSA subsampling.

## Data Availability

All prediction results to generate the figures are openly available in Zenodo at https://doi.org/10.5281/zenodo.17811671.

## Code Availability

The source codes of AF3-ReD and the AF3 MSA subsampling are available at https://github.com/OkazakiLab/af3_red and https://github.com/OkazakiLab/af3_mmm, respectively.

## Acknowledgements

All the structure predictions with AF3-ReD and the AF3 MSA subsampling were performed using the Research Center for Computational Science, Okazaki, Japan (Projects 22-IMS-C189, 23-IMS-C201, 24-IMS-C198, and 25-IMS-C227). This work was supported by JSPS KAKENHI Grant Numbers JP23K14160 (to J.O.), JP22H02595, JP23K23858, and JP25H02299 (to K.O.).

## Ethics declarations

### Competing interests

The authors declare no competing interests.

## Supplementary Information

The online version contains supplementary material available at…

## Notes

https://github.com/OkazakiLab/af3_red

https://github.com/OkazakiLab/af3_mmm

### Competing Interest Statement

The authors have declared no competing interest.

