## Supplementary Information for "Enhanced sampling of protein conformations in AlphaFold3 with repulsive bias in the diffusion generative model"

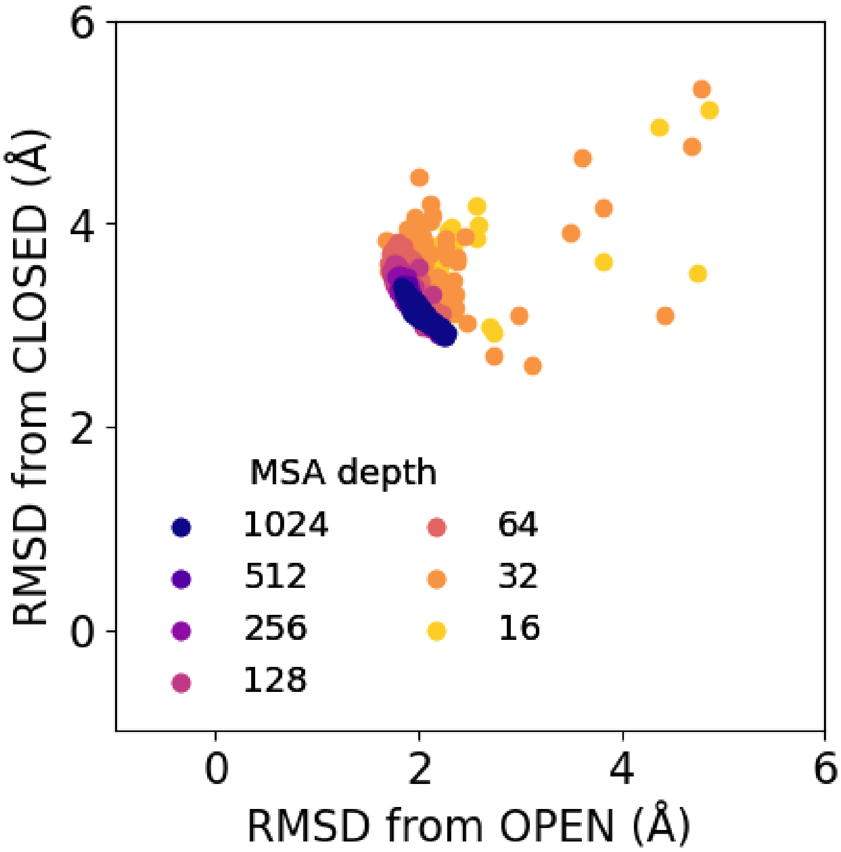


**Supplementary Fig. 1. Structure prediction of $F_1$-ATPase β subunit with AlphaFold2**

Root mean square deviations (RMSDs) of the structures predicted with MSA-subsampled AlphaFold2 for the apo $TF_1\beta$. RMSDs were calculated with respect to the open and closed structures extracted from the $TF_1$ ATPase complex determined by cryo-electron microscopy (PDB ID: 7XKH).

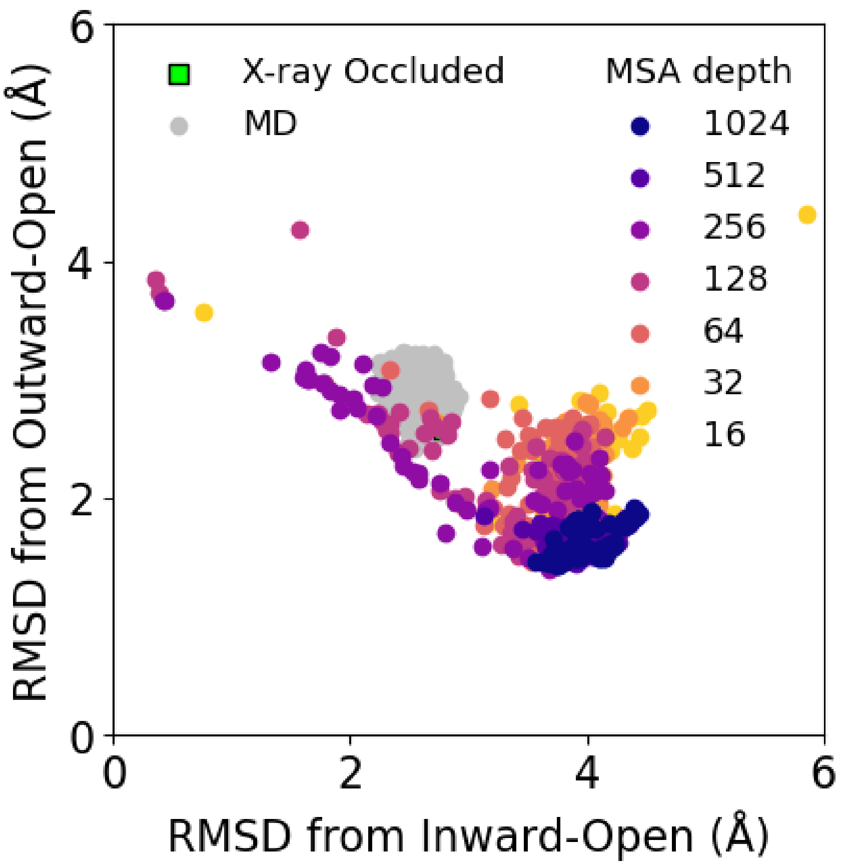


**Supplementary Fig. 2. Structure prediction of the oxalate-transporter OxlT with AlphaFold2**

RMSDs of the structures predicted with MSA-subsampled AlphaFold2 for the apo OxlT. RMSDs were calculated with respect to the outward-open crystal structure (PDB ID: 8HPJ) and the inward-open structure predicted with the original AF3. For the latter, the prediction with the highest pLDDT for the apo OxlT was used as the reference. The occluded crystal structure (green, PDB ID: 8HPK) and the MD simulation structures initiated from it (gray) are also plotted.

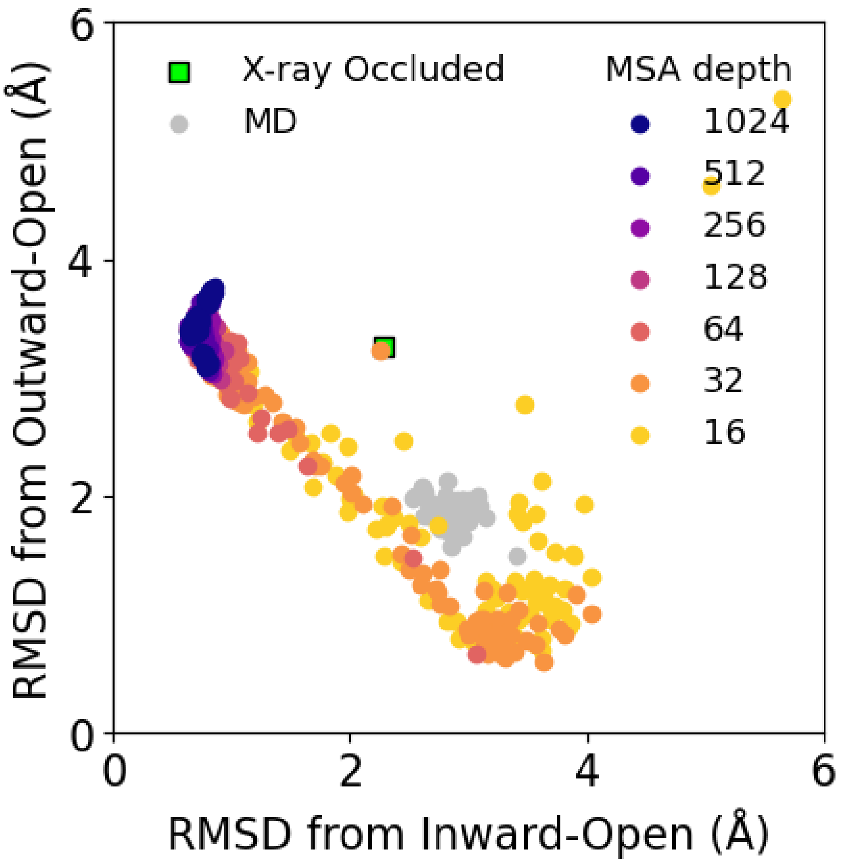


**Supplementary Fig. 3. Structure prediction of the nitrate-transporter NarK with AlphaFold2**

RMSDs of the structures predicted with MSA-subsampled AlphaFold2 for the apo NarK. RMSDs were calculated with respect to the inward-open crystal structure (PDB ID: 4U4V) and the outward-open structure predicted with the original AF3. For the latter, the prediction with the highest pLDDT for the apo NarK was used as the reference. The occluded structures from X-ray crystallography (green, PDB ID: 4U4W) and from our previous MD simulation (gray) are also plotted.

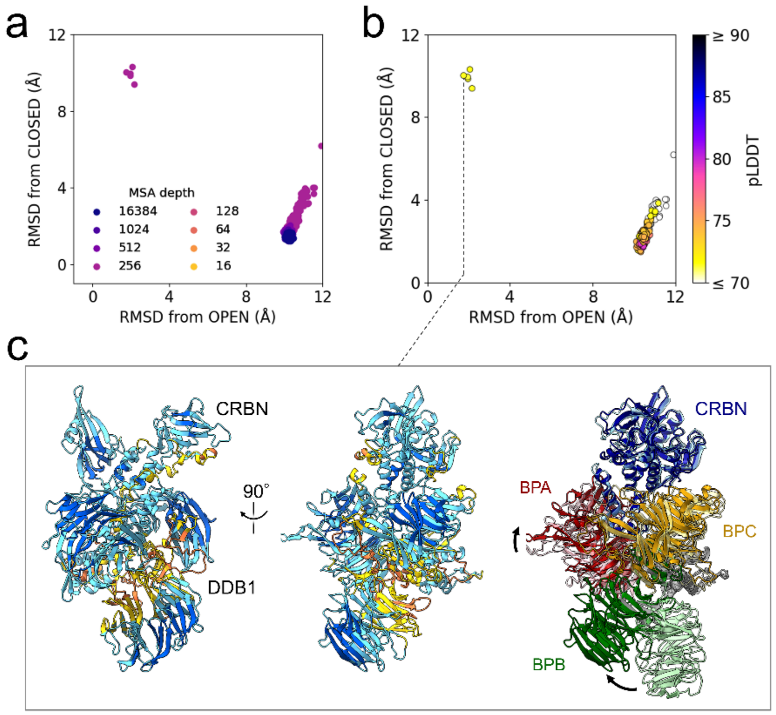


**Supplementary Fig. 4. the E3 ubiquitin ligase complex with AlphaFold3 MSA subsampling**

**a** RMSDs of the structures predicted with MSA-subsampled AlphaFold3 for the apo CRBN-DDB1. Note that for MSA depths lower than 256, the RMSDs are too large to be plotted within the display range of the figure. **b** RMSDs of the structures predicted with AF3 MSA subsampling at an MSA depth of 256. Points are colored according to the pLDDT score averaged over all atoms. **c** An open conformation predicted with AF3 MSA subsampling at an MSA depth of 256. The left and center panels show the pLDDT scores for each residue (blue: pLDDT ≥ 90; cyan: 70 ≤ pLDDT < 90; yellow: 50 ≤ pLDDT < 70; orange: pLDDT < 50). In the right panel, the predicted structure is superimposed on the CRBN of the crystal structure (shown in transparent colors), where CRBN and the three DDB1 domains (BPA, BPB, and BPC) are colored separately. The black arrows indicate the deviations of each DDB1 domain in the predicted structure from the crystal structure.
